# To Be or Not to Be an Oocyte: Msps/XMAP215 Controls Oocyte Cell Fate in the *Drosophila* Ovary

**DOI:** 10.64898/2026.02.10.705054

**Authors:** Wen Lu, Margot Lakonishok, Hannah Neiswender, Graydon B. Gonsalvez, Vladimir I. Gelfand

## Abstract

Cell fate determination, the process by which cells commit to specific identities, is fundamental to development. Oocyte specification in *Drosophila* provides an excellent model to dissect these molecular mechanisms. In the *Drosophila* ovary, 16 interconnected germline cells arise through four mitotic divisions with incomplete cytokinesis. Among them, the two oldest cells become pro-oocytes, and one is selected to develop into the oocyte while maintaining diploidy; the remaining 15 differentiate into nurse cells and enter endoreplication. Previously, we showed that Mini spindles (Msps), a microtubule polymerase and homolog of XMAP215, is essential for oocyte maintenance. Here, we report that Msps/XMAP215 is expressed early in development, with both mRNA and protein enriched in the pro-oocytes prior to oocyte specification. Knockdown of Msps prevents oocyte specification, leading to egg chambers with 16 nurse cells. Loss of Msps also disrupts the accumulation of the oocyte marker Orb and the microtubule minus-end binding protein Patronin/CAMSAP, both critical for oocyte specification. Remarkably, optogenetic recruitment of Msps is sufficient to increase microtubule polymerization and promote Orb accumulation in nurse cells, suggesting that Msps activity is sufficient to drive oocyte fate determination. Furthermore, we show that Msps associates with the germline-specific organelles, spectrosome and fusome, and becomes asymmetrically distributed among sister cells, allowing the two pro-oocytes to inherit higher levels of Msps than their siblings. Together, we propose a model in which Msps-mediated microtubule polymerization provides the pro-oocyte with a competitive advantage, initiating a positive feedback loop that involves dynein-dependent transport of *msps* mRNA to reinforce oocyte specification.

**Teaser:** A self-reinforcing cytoskeletal feedback loop selects a single egg cell from a cluster of interconnected germ cells.

## Introduction

In the *Drosophila* ovary, oocyte specification takes place within the anterior-most region of each ovariole, the germarium. The germarium is organized into several regions that support stepwise germ cell development. Germline stem cells (GSCs) are located at the anterior tip and divide asymmetrically to produce cystoblasts (CBs). Each CB undergoes four rounds of incomplete mitotic divisions, giving rise to 16 interconnected cystocytes. These divisions occur in region 1, while region 2a contains newly formed interconnected 16-cell cysts. As cysts progress into region 2b, they become lens-shaped and are enveloped by somatic follicle cells (Figure 1A) (*1*). In region 3, the 16-cell cyst becomes fully encapsulated by a monolayer of follicle cells and buds off as an individual egg chamber, marking the beginning of oogenesis in the vitellarium (stage 1).

**Figure 1.**
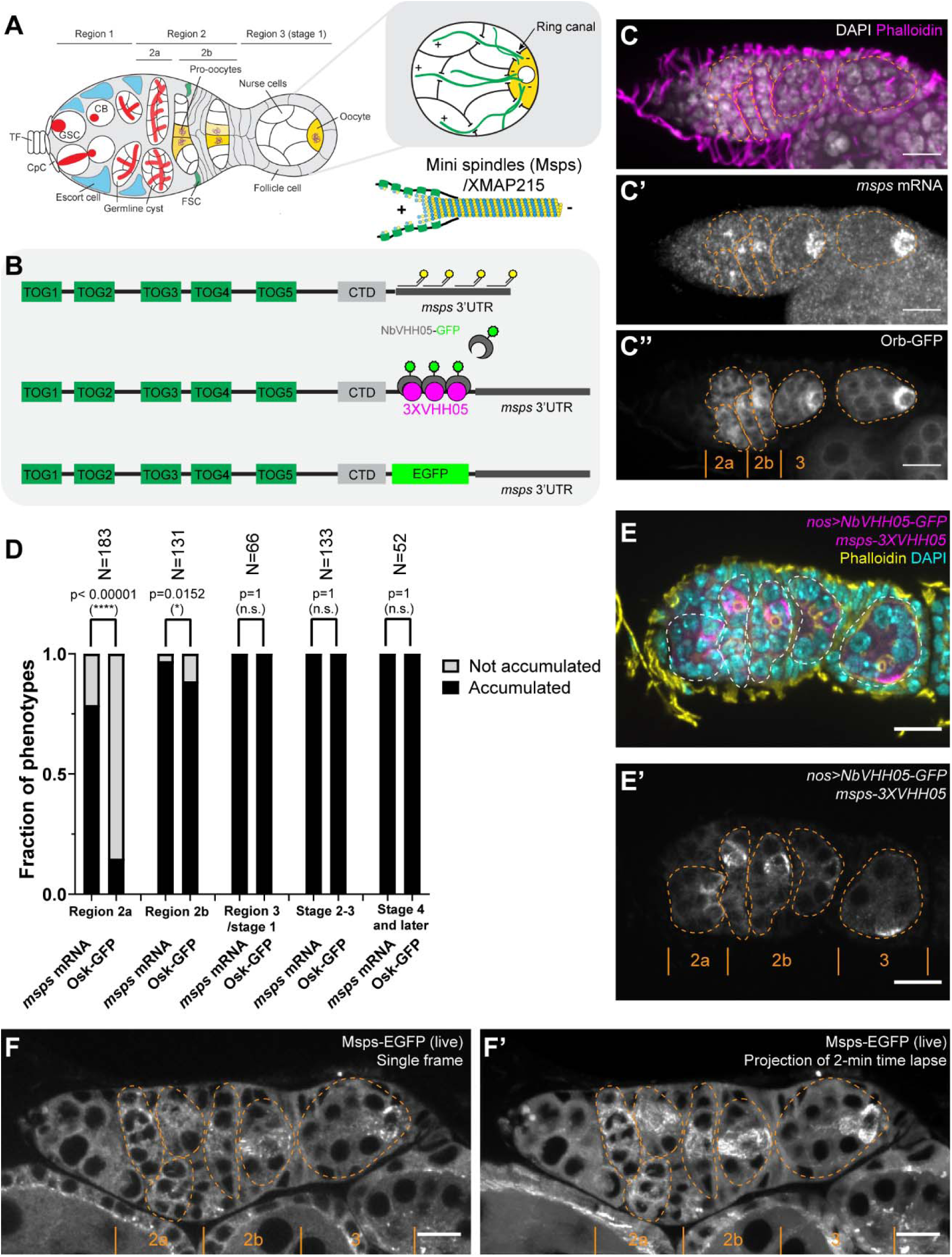
Msps is concentrated in pro-oocytes and early oocytes. (A) Left: schematic of a *Drosophila* germarium with spectrosome/fusome structures in red, synaptonemal complex in purple, and pro-oocytes/early oocytes in yellow. Inset: microtubule organization in early egg chambers, with most minus ends originating in the oocyte and plus ends extending through ring canals into nurse cells. Right: Mini spindles (Msps), the *Drosophila* homolog of XMAP215, promotes microtubule polymerization by delivering tubulin dimers to the plus end via its five TOG domains. (B) Domain organization of Msps/XMAP215 and methods used to visualize *msps* mRNA and protein localization. (C–C’’) Representative *msps* mRNA localization in an Orb-GFP germarium. *msps* mRNA was detected by smiFISH against the 3′UTR (C’). (D) Quantification of *msps* mRNA and Orb-GFP accumulation across germarium regions. Statistics between groups were performed using Fisher’s exact test. (E–E’) Representative Msps protein localization in a fixed germarium. Msps-3XVHH05 was detected by germline-specific expression of a GFP-tagged nanobody against VHH05 (driven by *nos-Gal4^[VP16]^*) (E’). (F–F’) Representative images of Msps-EGFP in a live germarium (single frame or 2-min time-lapse projection). See also Video 1. (C–C’’, E–F’) Orange dashed lines outline germline cysts. Scale bars, 10 µm.

The spectrosome, a spherical precursor of the fusome, is present in GSCs and CBs and helps to orient asymmetric divisions (*2*). During subsequent divisions, the spectrosome transforms into the branched fusome, an organelle enriched in membranous and cytoskeletal proteins. The fusome plays a critical role in anchoring the mitotic spindle during germ cell division, establishing cyst polarity and directing the asymmetric distribution of cell fate determinants, as well as organizing the microtubule network (*3, 4*). Although the fusome connects all 16 cystocytes through intercellular cytoplasmic bridges, known as ring canals, it is biased toward the two oldest cells, which inherit the most fusome material (*5*). This bias correlates with their entry into meiosis and pro-oocyte identity, marked by the appearance of C(3)G staining, a synaptonemal complex component (*6*). Ultimately, one of the pro-oocytes maintains meiosis and is selected as the oocyte, while the other exits meiosis and differentiates into a nurse cell, joining the other 14 cystocytes in endoreplication. Oocyte selection is marked by the enrichment of key molecular markers such as Orb, an RNA-binding protein essential for oocyte identity and axis specification (*7, 8*), as well as by the accumulation of centrosomes migrated from sister nurse cells (*9, 10*). Nurse cells then synthesize mRNAs and proteins, and deliver these materials to the developing oocyte through the ring canals to support rapid oocyte growth (*11–14*).

Oocyte specification depends on several molecular complexes that act in concert. (1) The fusome: loss of essential components such as the Adducin-like gene hu li tai shao (*hts*) (*15*) and α-Spectrin (*16*), disrupts fusome architecture and abolishes oocyte specification, yielding cysts with only nurse cells. (2) The microtubule network: disruption of the polarized microtubule network (*17*) or lack of its key regulators, such as microtubule minus-end binding protein Patronin/CAMSAP (*18*) and the microtubule-actin crosslinker Short stop (Shot) (*12, 19*), prevents oocyte specification. (3) Dynein-based transport: loss of dynein heavy chain (Dhc64C) (*20*), or its cofactors Bicaudal-D (BicD) (*21*), Egalitarian (Egl) (*22*), and Lis1 (*23*) results in cysts with 16 polyploid nurse cells and no diploid oocyte. (4) RNA-binding proteins: Orb is essential for oocyte specification (*7, 24*), functioning through translational activation of key oocyte transcripts such as *oskar* (*25*) and *gurken* (*26*). In summary, the fusome organizes a polarized microtubule network with the minus ends enriched in the future oocyte, enabling dynein-dependent transport of oocyte-specific factors to concentrate in this cell. This robust polarized transport along the microtubule network ensures the selection and maintenance of a single oocyte in each cyst. However, the upstream factor(s) that initiate microtubule polarization and thereby define the oocyte specification have remained unclear.

We previously demonstrated that the microtubule polymerase Mini Spindles (Msps), the *Drosophila* homolog of XMAP215, is essential for oocyte growth and oocyte fate maintenance (*27*). Msps/XMAP215 contains an N-terminal array of five conserved TOG (tumor overexpressed gene) domains (Figure 1B) (*28*). XMAP215/Msps tracks the microtubule plus-ends and enhances microtubule polymerization rate by increasing tubulin dimer concentration at the growing tips (*29, 30*). TOG1–2 engage free tubulin, and TOG5 associates with lattice-incorporated tubulin, while TOG3–4 function as a bridge that stabilizes the intermediate conformation between soluble and incorporated tubulin (Figure 1A–1B) (*31*). Loss of XMAP215/Msps disrupts spindle integrity, producing abnormally short or disorganized spindles during mitosis and meiosis in *Drosophila* (*32–34*). In the ovary, we showed that dynein-dependent transport of *msps* mRNA concentrates *msps* mRNA and Msps protein in the oocyte, where it stimulates microtubule polymerization and enhances nurse cell–to–oocyte transport, which is essential for oocyte fate maintenance (*27*).

In this study, we show that Msps/XMAP215 is not only essential for maintaining oocyte fate but also plays a key role in oocyte specification. We find that *msps* mRNA and Msps protein are enriched in pro-oocytes before oocyte selection. Loss of Msps disrupts oocyte specification, as evidenced by the failure to accumulate key oocyte determinants such as Orb and Patronin, resulting in egg chambers containing 16 nurse cells and no oocyte. Remarkably, optogenetic recruitment of Msps enhances microtubule polymerization and increases Orb level in nurse cells, indicating that Msps activity is sufficient to drive oocyte fate. Finally, we show that Msps associates with the spectrosome and fusome and is asymmetrically inherited by pro-oocytes, supporting a model in which localized Msps-stimulated microtubule growth thus provides a competitive advantage to one cell and initiates a positive feedback loop to reinforce oocyte specification.

## Results

### Msps is concentrated in pro-oocytes and early oocytes

A polarized microtubule network is essential for oocyte specification in the *Drosophila* germarium (*11*). We, therefore, examined the localization of the microtubule polymerase Msps in the germarium. Using single-molecule inexpensive fluorescence *in situ* hybridization (smiFISH) (Figure 1B), we found that *msps* mRNA is enriched in discrete foci beginning in region 2a, preceding detectable accumulation of the oocyte marker Orb (Figure 1C–1D). This suggests that *msps* mRNA enrichment marks an early step in oocyte specification, before Orb-dependent mechanisms are established.

To analyze protein distribution, we used two CRISPR knock-in lines: an indirect labeling by the NanoTag VHH05 that can be recognized by EGFP-tagged nanobody (Msps-3XVHH05 with nanobody-EGFP) (*27*) and a direct C-terminal EGFP fusion (Msps-EGFP). Both alleles are homozygous viable and fertile, indicating that the tags do not impair protein function. In both cases, Msps protein was enriched in pro-oocytes from region 2a and became progressively concentrated in the oocyte in regions 2b and 3 (Figure 1E–1F′; Video 1). Co-labeling with the synaptonemal complex marker C(3)G confirmed that Msps-EGFP accumulated in pro-oocytes undergoing meiosis in region 2a (Supplementary Figure 1A–1B). Notably, in some cysts, both pro-oocytes have entered meiosis, yet only one showed strong Msps enrichment (Supplementary Figure 1C–1D), suggesting a transitional phase in which Msps accumulation precedes the eventual selection of a single oocyte.

In summary, both *msps* mRNA and Msps protein are enriched in pro-oocytes and early oocytes, indicating that Msps may act as an early determinant of oocyte specification.

### Msps is required for oocyte specification

As Msps is essential for spindle assembly and mitotic progression (*32*), early germline knockdown using the strong driver *nanos-Gal4[VP16]* (*35, 36*) eliminates the germline entirely (*27*). To bypass this germless phenotype, we employed a weaker *nanos-Gal4* line (lacking the VP16 activation domain) in combination with the temperature-sensitive *tubP-Gal80[ts]* (*37, 38*). At 18°C, Gal80[ts] suppresses *msps-RNAi* expression in germline cells (RNAi OFF), whereas at 30°C Gal80[ts] is inactivated, allowing germline-specific *msps-RNAi* expression (RNAi ON) (Figure 2A).

**Figure 2.**
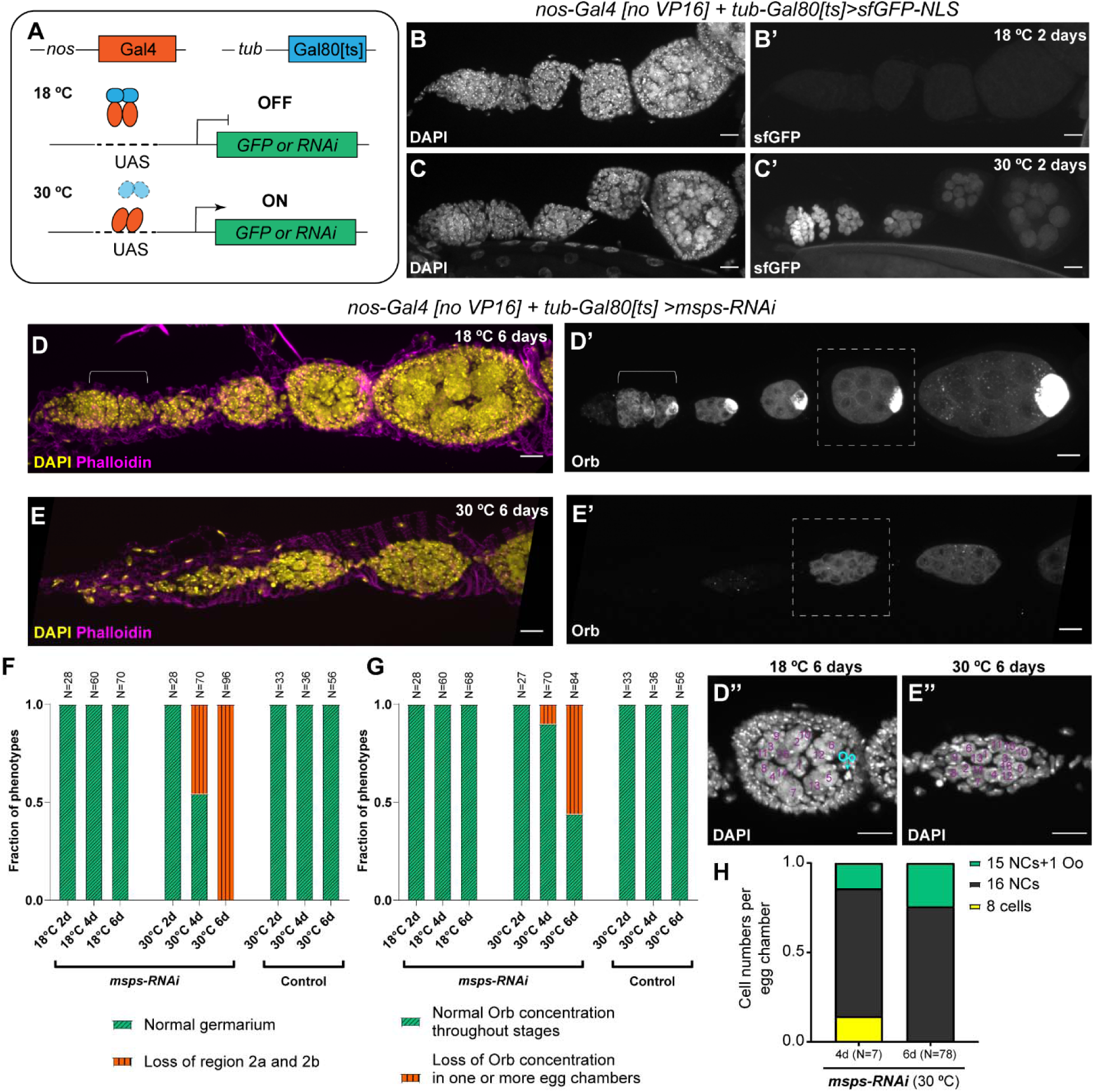
Msps is required for oocyte specification. (A) Schematic of the temperature-sensitive Gal80 system. At the permissive temperature (18 °C), Gal80[ts] binds Gal4 and prevents its interaction with upstreaming activating sequences (UAS), repressing transgene expression (OFF). At the restrictive temperature (30 °C), Gal80[ts] is inactivated, allowing Gal4 to activate UAS-driven transgenes (ON). (B–C’) Germline expression of nuclear-localized superfolder GFP (sfGFP-NLS) using the Gal80[ts] system. Flies were raised at 18 °C, and F1 progeny were either maintained at 18 °C or shifted to 30 °C for 2 days before examination. Maximal z-projections of the GFP signal are shown in both cases. (D–E’) Germline expression of *msps*-RNAi using the Gal80[ts] system. Flies were raised at 18 °C, and F1 progeny were either maintained at 18 °C or shifted to 30 °C for 6 days before examination. (D-D’) The white brackets indicate region 2a–2b of the germarium. (D’’–E’’) DAPI staining of the white dashed boxes in (D–E). (F–G) Quantification of germaria lacking region 2a/2b and of oocyte specification defects in the listed genotypes. All crosses were raised at 18 °C; F1 progeny were maintained at 18 °C or shifted to 30 °C for the indicated number of days. Sibling flies carrying *tub-Gal80[ts]* but lacking *msps*-RNAi served as controls. (H) Summary of cell numbers in egg chambers lacking Orb concentration. (B–E’’) Scale bars, 10 µm.

We first validated the efficiency of the temperature-sensitive Gal80[ts] system using a nuclear GFP reporter (*UASp-sfGFP-NLS*). Shifting flies from 18°C to 30°C was sufficient to induce robust GFP expression in germline cells (Figure 2B–2C′; Supplementary Figure 2A–2B).

Having validated that the temperature-sensitive Gal80[ts] system efficiently induces transgene expression upon temperature shift, we next used it to induce the expression of *msps-RNAi*. Upon shifting adult females from 18°C to 30°C, we observed progressive defects. After 4–6 days, many ovarioles lost their germaria due to impaired germline cell divisions (Figure 2D–2F), while the egg chambers that formed showed striking specification defects. In these chambers, Orb frequently failed to accumulate, and a diploid oocyte nucleus was not detected (Figure 2D–2G). By contrast, flies carrying *tub-Gal80[ts]* but lacking *msps-RNAi* showed no detectable defects in germline divisions or oocyte specification after 6 days at 30 °C (Figure 2F-2G), confirming that the observed phenotypes are due to loss of *msps*. Most *msps-RNAi* affected chambers contained 16 polyploid nurse cells, consistent with a complete failure of oocyte specification rather than defective cyst formation (Figure 2D’’-2E’’ and 2H; Supplementary Figure 2C-2F; Videos 2–3). A smaller fraction contained 15 polyploid nurse cells and a single diploid cell without Orb enrichment (Figure 2H), suggesting additional defects in oocyte fate maintenance, consistent with our previous findings (*27*).

Together, these results demonstrate that Msps is not only essential for germline divisions and oocyte maintenance, but also required for the initial specification of the oocyte in the *Drosophila* ovary.

### Msps and Patronin are interdependent in oocyte localization

The microtubule minus-end–binding protein Patronin, the sole *Drosophila* member of the CAMSAP family, is required for symmetry breaking in pro-oocytes and thus for oocyte specification (*18*). To test whether Msps and Patronin act together, we first examined their localization simultaneously. smiFISH staining of *msps* mRNA in ovaries expressing endogenously tagged Patronin-YFP (Nashchekin et al., 2016) revealed co-accumulation of *msps* mRNA and Patronin protein as early as in region 2a, and it persisted throughout the germarium (Figure 3A–A′′). Similarly, Msps-EGFP in ovaries with endogenously tagged Patronin-mKate (*18*) showed Msps-EGFP comets emerging from Patronin-mKate foci in the earliest oocytes (Figure 3B-3C’’; Video 4). These results suggest that Msps and Patronin function in concert during oocyte specification.

**Figure 3.**
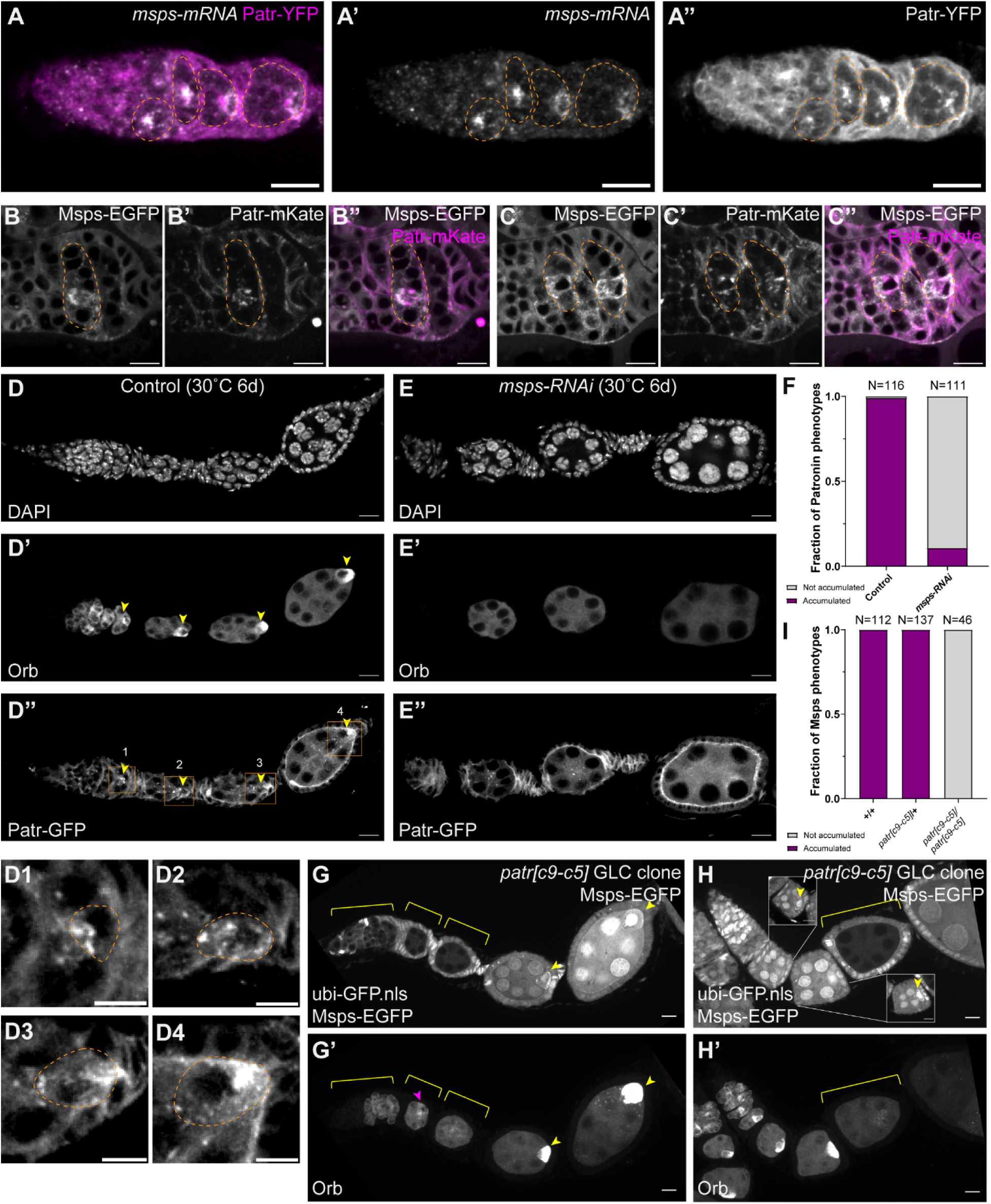
Msps and Patronin are interdependent for oocyte specification and localization. (A–A’’) Localization of *msps* mRNA (smiFISH) and YFP-tagged Patronin protein. Orange dashed lines, germline cysts. (B–C’’) Localization of EGFP-tagged Msps and mKate-tagged Patronin. Maximal projection of 2-min time-lapse live imaging at two focal planes of the same germarium (see also Video 4). Orange dashed lines, germline cysts. (D–E’’) Orb staining (D’–E’) and Patronin localization (*ubi-Patronin-GFP*, D’’–E’’) in control (D–D’’) and *msps*-RNAi (E–E’’) samples. Flies were raised at 18 °C and shifted to 30 °C for 6 days. Accumulation of Orb and Patronin in the oocyte (yellow arrowheads, D’–D’’; D1-D4, zoom-in areas of the regions indicated by the orange boxes in D’’, with oocytes outlined by orange dashed lines) was abolished in *msps*-RNAi (E’–E’’). (F) Quantification of Patronin localization phenotypes in control and *msps*-RNAi samples. (G–H’) Msps-EGFP (G–H) and Orb staining (G–H’) in *patr[c9-c5]* FRT clones. Homozygous clones (yellow brackets) lack nuclear GFP; heterozygous cells retain nuclear GFP. Yellow arrowheads mark Msps-EGFP and Orb accumulation in heterozygous oocytes. Insets in (H) show Msps-EGFP accumulation at different focal planes. Magenta arrowhead in (G’) highlights residual Orb signal in *patr[c9-c5]* homozygous clones. (I) Quantification of Msps localization phenotypes in control clones (FRT G13), *patr[c9-c5]* heterozygotes, and *patr[c9-c5]* homozygous clones. (A–C’’, D–E’’, G–H’) Scale bars, 10 µm. (D1-D4) Scale bars, 5 µm.

To test their interdependence, we perturbed each protein and examined the localization of the other. In *msps-RNAi* ovaries, Patronin-GFP failed to accumulate, accompanied by loss of oocyte specification, as indicated by absent Orb enrichment and the lack of a diploid oocyte nucleus (Figure 3D–3F). Conversely, in *patronin* null mutant clones (*patr[c9-c5]*), oocyte specification was disrupted, consistent with the previous report from the St Johnston’s lab (*18*), and Msps-EGFP accumulation was lost in homozygous clones but maintained in heterozygous and control clones (Figure 3G–3I; Supplementary Figure 3A–3D).

Finally, genetic interaction further supported this functional link. Having one endogenous copy of Msps tagged with EGFP partially rescued the oocyte specification defects in *patronin* null clones, as indicated by Orb enrichment (magenta arrowheads, Figure 3G’ and Supplementary Figure 3C–3D). By contrast, adding an extra copy of GFP-tagged Patronin (*ubi-Patronin-GFP*) enhanced the severity of *msps*-RNAi phenotypes (Supplementary Figure 3E). These results show that altering the dosage or stability of one component changes the phenotypic penetrance of the other, underscoring their functional interdependence.

Together, these results indicate that Msps and Patronin are interdependent for both oocyte specification and localization. However, it remains unclear whether this reflects a direct protein-protein interaction or an indirect consequence of failed oocyte selection.

### Msps is sufficient to promote oocyte specification

Having shown that Msps and Patronin are interdependent, we next asked which of them acts upstream. In *msps*-RNAi ovaries, we occasionally observed Patronin accumulation in germline cells that nevertheless lacked Orb enrichment and a diploid oocyte nucleus carrying karosome formation (magenta arrowheads, Supplementary Figure 3F–F′′), suggesting that Patronin alone is insufficient to specify an oocyte without Msps. We therefore tested whether ectopic accumulation of Msps is sufficient to drive oocyte specification.

Oocytes normally exhibit higher microtubule polymerization activity than their sibling nurse cells (*12, 18*). This activity is dependent on Msps accumulation in the oocyte (*27*). Elevated microtubule polymerization activity in the oocyte establishes a polarized microtubule network, with minus-ends concentrated in the oocyte and plus-ends extending into nurse cells through the ring canals, thereby enabling dynein to transport and concentrate oocyte determinants to the oocyte. We reasoned that ectopic recruitment of Msps in nurse cells would increase local microtubule growth, potentially altering this microtubule polarity and concentrating oocyte fate determinants outside of the oocyte.

To achieve this, we developed an optogenetic system, OptoMsps, to recruit Msps onto microtubules (Figure 4A). This strategy combines the improved light-inducible dimer (iLID) system (*39*) with the NanoTag–nanobody system (*40*). The iLID module consists of an *Avena sativa* LOV2 domain fused to an SsrA peptide, which is sterically blocked in the dark but uncaged by blue light, enabling high-affinity binding to SspB. For microtubule anchoring, we adapted a component of the Opto-Katanin system (*41*), using the N-terminal microtubule-binding domain of EB3—homologous to the *Drosophila* EB-SUN protein (*42*)—fused to the blue light–sensitive dimerization module VVDfast (*43*), followed by the iLID (LOV2-SsrA) domain. Upon blue light exposure, EB3N rapidly dimerizes, increasing its affinity for microtubules, while the SspB–nanobody fusion recruits endogenously tagged Msps to microtubules via the interaction between SsrA and SspB. In this way, local illumination recruits Msps outside of the oocyte and into neighboring nurse cells, where it can enhance microtubule polymerization (Figure 4B).

**Figure 4.**
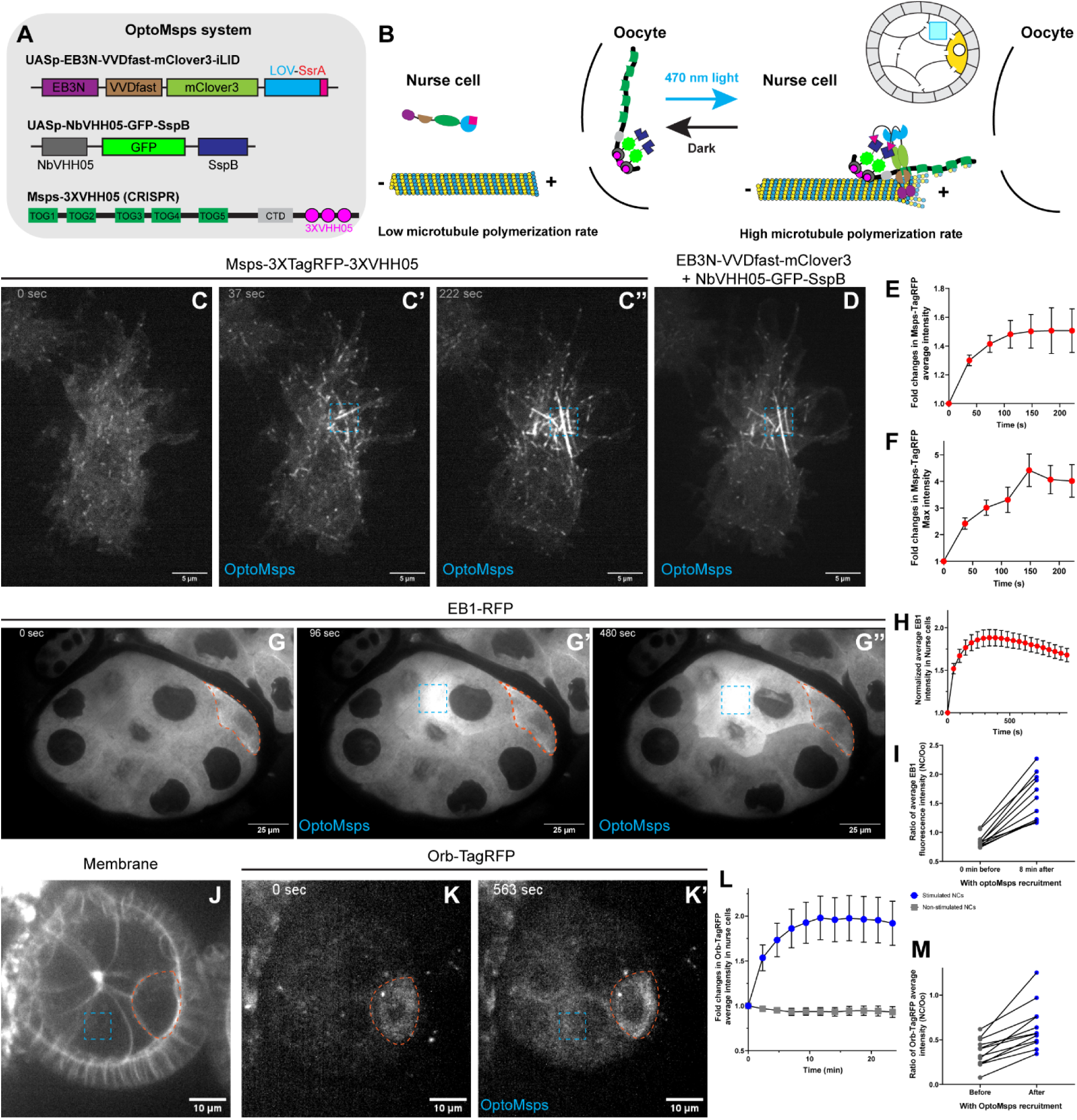
Msps is sufficient to promote oocyte specification. (A) Schematic of the OptoMsps system, composed of three elements: (1) a transgenic microtubule-binding domain of EB3 (EB3N) fused to the blue light–sensitive dimerization module VVDfast, followed by LOV2-SsrA; (2) a transgenic fusion of SspB with a nanobody recognizing the VHH05 NanoTag; and (3) a CRISPR knock-in of three VHH05 NanoTags into the endogenous *msps* locus (Msps-3XVHH05). (B) Cartoon illustrating OptoMsps recruitment: blue light induces VVDfast homodimerization and LOV2-SsrA/SspB heterodimerization, driving the recruitment of Msps to microtubules. Local stimulation in nurse cells artificially increases microtubule polymerization activity. (C–F) Validation of OptoMsps in *Drosophila* S2R+ cells. (C–C’’) Msps-3XTagRFP-3XVHH05 before and after local blue-light illumination (blue dashed box). (D) End-point GFP channel showing localization of the two OptoMsps components. (E–F) Quantification (mean ± SEM, N=7) of normalized average and maximum TagRFP intensity in the illuminated region. See also Video 5. (G–I) OptoMsps increases EB1 signal in *Drosophila* nurse cells. (G–G’’) EB1-RFP before and after local blue-light illumination in nurse cells (blue dashed box). Orange dashed outline, oocyte. (H) Quantification (mean ± SEM, N=12) of normalized average EB1-RFP intensity in the entire illuminated nurse cells. (I) Ratio of EB1-RFP intensity between illuminated nurse cells and the oocyte before and 8 min after illumination (N=11). See also Video 7. (J–M) OptoMsps increases Orb intensity in *Drosophila* nurse cells. (J) Membrane outline of the egg chamber, with oocyte marked (orange dashed line) and blue-light illumination region (blue dashed box). (K–K’’) Orb-TagRFP before and after local Msps recruitment in the nurse cells. (L) Quantification of normalized average Orb-TagRFP intensity in nurse cells (blue dots, N=12) versus non-illuminated nurse cells in neighboring chambers (gray dots, N=13). (M) Ratio of Orb-TagRFP intensity between nurse cells and the oocyte before and after illumination (N=12). See also Video 7.

We first validated OptoMsps in *Drosophila* S2R+ cells. Local blue light stimulation induced rapid accumulation of Msps-3XTagRFP-3XVHH05 on microtubules (Figure 4C–4F; Video 5), confirming efficient recruitment by the OptoMsps system. Notably, the recruited signal often spread beyond the original illuminated region (Figure 4C-4D), likely due to light scattering and the intrinsic ability of Msps to track growing microtubule ends once recruited.

We next tested OptoMsps in egg chambers. Global illumination showed rapid recruitment of the OptoMsps components to microtubules within a couple of seconds (Supplementary Figure 4A–A’’; Video 6). To measure functional consequences of Msps recruitment, we used EB1-RFP as a readout of microtubule polymerization activity. Although individual EB1 comets could not be resolved under optogenetic imaging conditions, local Msps recruitment consistently increased EB1-RFP signal in the illuminated area (Figure 4G–G′′; Video 7). Due to the geometry of the nurse cells and the spreading nature of the OptoMsps, the recruitment typically involved two to three nurse cells rather than a single target cell. Nevertheless, blue light stimulation consistently produced a clear increase in EB1-RFP signal, which subsequently spread throughout the entire nurse cell cytoplasm (Figure 4H–4I).

Finally, we asked whether Msps recruitment could alter oocyte fate markers. Using a MiMIC-based system (*44*), we generated an endogenous tagged Orb-TagRFP line (Venken et al., 2011), which completely recapitulates Orb antibody staining and fully functional as it supports normal oocyte specification when *in trans* with the *orb* null allele (Supplementary Figure 4B–4D”). Using this reporter, we found that local Msps recruitment in nurse cells induced a consistent increase in Orb signal, whereas unstimulated nurse cells showed no change over time (Figure 4J–4L; Video 8). Quantification confirmed a consistent post-stimulation increase of Orb-TagRFP in OptoMsps recruited nurse cells (Figure 4M).

Together, these results demonstrate that targeted recruitment of Msps is sufficient to enhance local microtubule polymerization and promote Orb accumulation in nurse cells, indicating that Msps activity alone can drive oocyte specification.

### Asymmetric segregation of Msps protein and dynein-dependent transport of its mRNA

Having established that Msps is both necessary and sufficient for oocyte specification, we next asked how the initial Msps asymmetry arises between the pro-oocytes and how it is propagated to a definitive fate decision.

Using two CRISPR knock-in lines (Msps-3XVHH05 and Msps-EGFP), we observed Msps association with the spectrosome and fusome in GSCs, CBs, and early cysts (Figure 5A; Supplementary Figure 5A). In later cysts, Msps appeared diffusely enriched around the branched fusome (Figure 5A; Supplementary Figure 5B). From region 2b to region 3, Msps concentrated either in both pro-oocytes or in the presumptive oocyte (Supplementary Figure 5C–5E), coinciding with the loss of clear fusome association. Since pro-oocytes are known to inherit more fusome material than their siblings (*5, 45*), it is likely that the spectrosome and fusome association biases Msps segregation toward the pro-oocytes. Supporting this idea, the spectrosome/fusome association of Msps was completely lost when the essential fusome component Hts was depleted by RNAi (Supplementary Figure 5F-5H). These findings suggest that Msps asymmetry in cystocytes is established, at least in part, through its association with the spectrosome and fusome structures.

**Figure 5.**
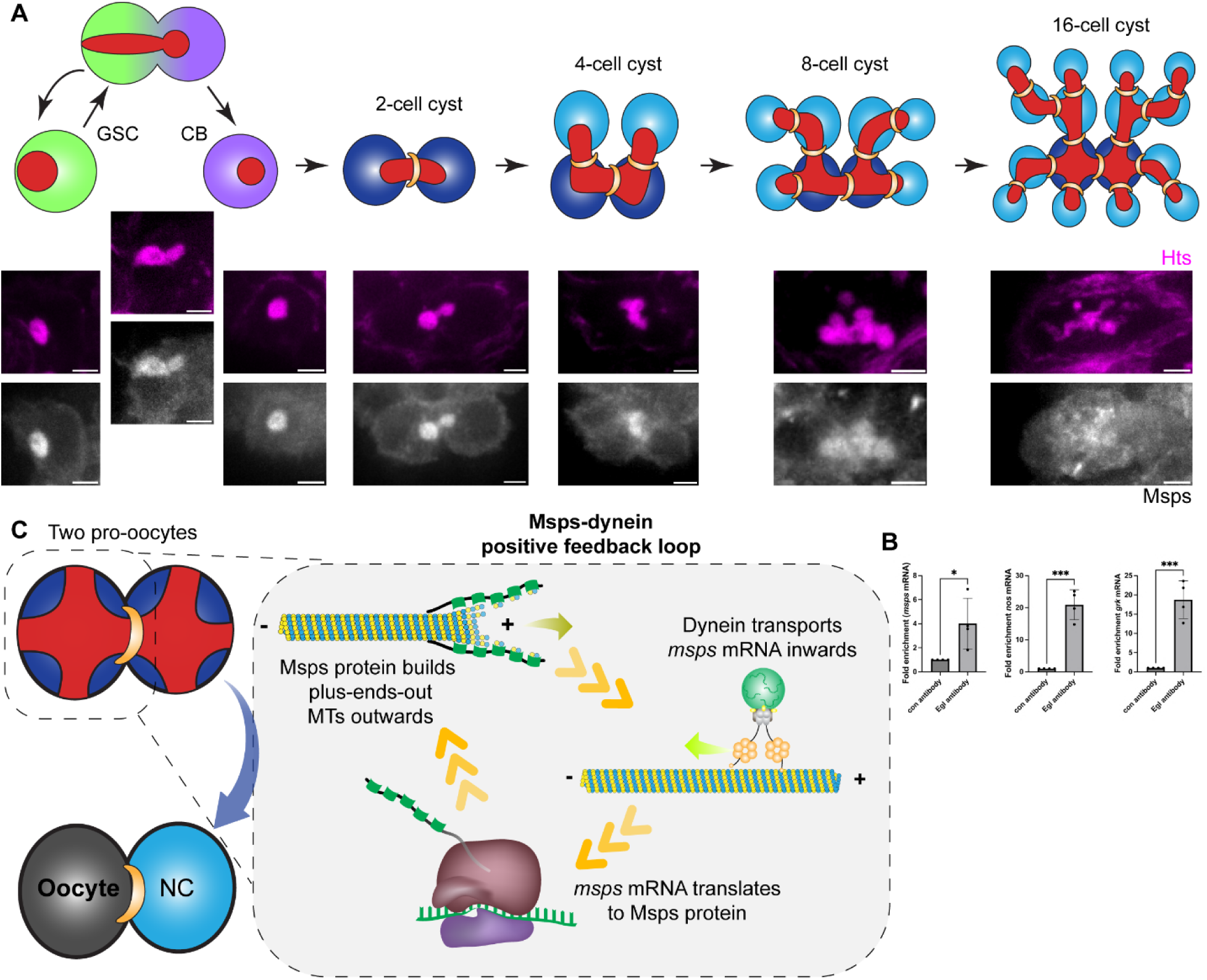
Msps initiates a positive feedback loop for oocyte specification. (A) Msps association with spectrosome and fusome structures in successive germline stages: germline stem cells (GSCs), cystoblasts (CBs), 2-cell, 4-cell, and 8-cell cysts. In 16-cell cysts, Msps shows a more diffuse “cloudy” distribution around the branched fusome. Scale bars, 2 µm. (B) Interaction of Egalitarian (Egl) with *msps* mRNA compared to two known Egl-dependent cargos, *nanos* (*nos*) and *gurken* (*grk*). (C) A positive feedback mechanism amplifies a small difference in Msps protein distribution between the two pro-oocytes into a definitive oocyte fate decision.

In addition to this protein-level segregation, we also asked whether *msps* mRNA is actively targeted to the oocyte by dynein-mediated transport. Previously, we have shown that *msps* mRNA accumulation in the oocyte requires dynein and the mRNA-binding adaptor Egalitarian (Egl) (*27*). Egl functions as the adaptor linking specific mRNAs to BicD and the dynein motor (*46–48*). Here, we examined the interaction between *msps* mRNA and cytoplasmic dynein complex. Immunoprecipitation of ovarian lysate with Egl antibody showed enrichment of *msps* mRNA compared to control (Figure 5B), indicating that *msps* mRNA specifically associates with the dynein complex via Egl. Together with our previous observation that inhibition of dynein or Egl disrupts *msps* mRNA accumulation in the oocyte (*27*), the present finding that *msps* mRNA interacts with Egl demonstrates that *msps* mRNA is a bona fide dynein cargo.

## Discussion

In this study, we demonstrate that the conserved microtubule polymerase Msps/XMAP215 is both necessary and sufficient for oocyte specification in *Drosophila*. Msps accumulates in pro-oocytes and early oocytes (Figure 1), and its loss abolishes oocyte specification (Figure 2). Msps and the minus-end binding protein Patronin are interdependent for oocyte localization and specification (Figure 3). Using optogenetic recruitment, we further show that increasing Msps activity outside the oocyte is sufficient to induce accumulation of oocyte fate markers (Figure 4). Finally, we find that Msps protein segregates with the fusome and that *msps* mRNA is transported by the dynein motor complex, allowing asymmetric enrichment in pro-oocytes (Figure 5).

Together, these findings support a positive feedback model in which Msps and dynein cooperate to amplify small initial asymmetries into a definitive oocyte fate. In this model: (1) the future oocyte inherits slightly more Msps than its sibling pro-oocyte, leading to increased microtubule growth with plus-ends out; (2) dynein transports more *msps* mRNA into this cell; and (3) more translated Msps protein further promotes more polarized microtubule assembly, which in turn drives even more dynein-dependent *msps* mRNA transport into the future oocyte. Through the cycles, an initial small bias in Msps is progressively amplified into a robust oocyte specification decision (Figure 5C).

### Oocyte selection: ‘predetermined’ or ‘stochastic’?

Two prevailing models have been proposed to explain how oocytes are selected. The first ‘predetermined’ model is that the oocyte is specified at the time of the first mitotic division of the cystoblast (in one of the pro-oocytes), and this asymmetry is maintained in subsequent development to a final oocyte selection. The second ‘stochastic’ model assumes that cell competition between the two pro-oocytes results in random selection of the future oocyte by the accumulation of oocyte-specific mRNAs, proteins, and/or organelles (*11*). For both models, the inheritance of fusome materials is the key to the establishment of asymmetry. A later paper from the Stanislav Shvartsman group carefully examined the fusome topology during development, and they found that fusome asymmetry between the pro-oocytes decreases with each cell division, while asymmetry between the pro-oocytes and the rest of the cyst increases (*5*). Thus, they propose the model of ‘equivalency with a bias’: the amplified fusome asymmetry between the pro-oocytes and the other sister cells chooses these two cells as candidates for oocyte selection; decreasing fusome asymmetry between the two pro-oocytes suggests that the oocyte selection between pro-oocytes is likely to be stochastic (*5*).

Our data best fit an ‘equivalency with a bias’ model, where fusome inheritance ensures both pro-oocytes are candidates, but Msps-dynein feedback resolves the competition stochastically. Msps, as a critical oocyte-selection factor, is not simply predetermined or stochastic. Because the fusome extends through intercellular bridges and is partitioned unequally during successive divisions, pro-oocytes inherit more fusome material than their sibling cystocytes (Figure 5A). Through the Msps-fusome association, they also reliably inherit more Msps, making them the primary candidates for oocyte fate. Subsequently, a small stochastic difference in Msps levels between the two pro-oocytes is sufficient to engage the positive feedback loop that drives a definitive oocyte fate decision. Furthermore, we noticed that Msps is cloudy around fusome at the later cyst stages and is not clearly associated with fusome in the region 2b pro-oocytes (Supplementary Figure 5), indicating that a stochastic difference may occur around later cyst stages to region 2a.

### Oocyte selection: flexible yet robust

Our data showed that *msps* mRNA is a cargo for the dynein motor complex. It binds to dynein through the interaction with the RNA-binding protein Egl and accumulates in the oocyte (Figure 5) (*27*). Intriguingly, the interaction between *msps* mRNA and Egl is less robust compared to other dynein mRNA cargoes, such as *gurken (grk)* and *nanos (nos)*. One contributing factor may be stage specificity: whereas *grk* and *nos* remain dynein cargos throughout oogenesis, *msps* mRNA functions mainly in early-to-mid stages for oocyte specification and fate maintenance. Because our immunoprecipitation assays used whole-ovary lysates, in which late-stage egg chamber materials are overrepresented, the developmental timing of *msps* transport may reduce the apparent strength of the *msps*–Egl interaction compared to *grk* or *nos*.

In addition, the relatively weak interaction may serve a functional purpose. The development proceeds relatively slowly in the germarium, where ∼4 days are required to reach region 3/stage 1, compared to the ∼3–4 days needed for the subsequent maturation of egg chambers through stages 1–14 (*49*). We propose that the relatively weak interaction between *msps* mRNA and the dynein motor complex allows Msps accumulation to build gradually, slowing the amplification of the positive feedback loop. This delay ensures that the commitment to oocyte fate is not made prematurely but instead occurs at the appropriate developmental stage in the germarium.

Furthermore, oocyte selection in the *Drosophila* germarium is not irreversible; the initially specified oocyte can revert to a nurse-cell fate if polarity cues, the microtubule network, or dynein-dependent transport are disrupted, underscoring the plasticity of the oocyte selection process (*50, 51*). We propose that a weaker *msps*-dynein interaction may contribute to this plasticity, permitting error correction before a definitive oocyte fate is established.

Interestingly, unlike most other Egl-dependent mRNAs, which typically carry a single RNA stem-loop localization signal (e.g., TLS in *K10*, GLS in *gurken*, ILS in the *I-factor* retrotransposon, and SL1 in *hairy*) (*46, 52–54*), *msps* mRNA contains multiple dynein localization elements distributed in both its coding sequence and 3′UTR (*27*). We propose that this unusual redundancy enhances the robustness of *msps* transport into the oocyte, ensuring reliable activation of the positive feedback loop and enforcing the specification of only one oocyte per cyst.

Positive feedback loops are a recurring theme in oocyte specification. Orb, the *Drosophila* homolog of Cytoplasmic Polyadenylation Element-Binding protein (CPEB), binds to cytoplasmic polyadenylation elements in its own 3′UTR to promote translation of *orb* mRNA, thereby reinforcing Orb accumulation specifically in the oocyte (*24, 55*). This self-sustaining mechanism ensures that once Orb begins to enrich in one cell, it is stabilized and amplified to consolidate oocyte fate. Notably, Orb accumulation becomes predominant in pro-oocytes only in region 2b, whereas *msps* mRNA and protein are already enriched in region 2a (Figure 1D and Supplementary Figure 1B). We therefore speculate that the Msps–dynein positive feedback loop acts earlier, followed by the Orb–*orb* positive feedback loop, and that together these mechanisms reinforce the robustness of oocyte specification to ensure that a single cell is selected from each cyst.

### Oocyte selection: Microtubule plus- and minus-end regulation

Microtubule organization is central to oocyte selection, requiring a coordinated balance between polymerization at plus ends and stabilization at minus ends. Msps/XMAP215, a conserved microtubule polymerase, promotes plus-end growth by accelerating tubulin addition, while Patronin/CAMSAP stabilizes microtubule minus ends against depolymerization. It is therefore not surprising that both proteins are required and interdependent for oocyte specification, as each contributes to establishing the polarized microtubule network that is required for dynein-mediated transport of oocyte determinants.

Intriguingly, our genetic data reveal a functional interaction between Msps and Patronin. Tagging one endogenous copy of Msps with EGFP partially rescues the oocyte specification defects in *patronin* null clones, as indicated by Orb accumulation. In contrast, introducing an extra copy of GFP-tagged Patronin driven by a weak ubiquitous promoter (*ubi-Patronin-GFP*) further enhances the defects caused by *msps*-RNAi (Supplementary Figure 3). These findings suggest that Msps and Patronin activities must be carefully balanced: excessive Patronin may stabilize minus ends without sufficient plus-end growth, whereas reduced Patronin function can be partly compensated by tuning Msps levels. Moreover, our findings highlight the need for caution, as tagging proteins with bulky fluorophores such as GFP can alter protein stability or affect protein–protein interactions. CRISPR-based tagging with smaller epitopes or nanotags may help minimize these potential artifacts. Together, these observations highlight the importance of coordinated regulation at both microtubule ends to ensure robust oocyte specification.

### Oocyte selection: conserved principles, divergent structures

Oocyte specification relies on conserved principles of cytoskeletal polarization and the selective accumulation of cell fate determinants; yet, the molecular machinery guiding this process exhibits both conservation and divergence across species. In *Drosophila*, the Msps/XMAP215–dynein positive feedback loop is embedded within the unique context of the spectrosome and fusome, organelles unique to insects and other invertebrates that organize the polarized microtubule network, and bias material transfer into the pro-oocytes. By contrast, most vertebrate germline cysts lack fusomes and instead rely on TEX14-stabilized intercellular bridges (*56*).

Despite these structural differences, the overall logic is similar: germline cells undergo incomplete cytokinesis and remain interconnected by intercellular bridges, through which sister cells transfer cytoplasmic materials to a single dominant oocyte (*57*). In flies, this is mediated by fusomes and ring canals, whereas in mammals, TEX14-stabilized intercellular bridges perform this role. Recent work has highlighted that centralspindlin and Ect2, core components required for stable intercellular bridges, are evolutionarily conserved across Metazoa, underscoring their dual importance in the emergence of multicellularity and in the maintenance of germline cyst connectivity (*58*). In both fly and mammalian systems, “nurse-like” cells donate cytoplasmic contents, such as mitochondria, Golgi, RNAs, and other organelles, via a microtubule-dependent manner, to support the future oocyte, before undergoing programmed cell death (*12, 13, 59–63*).

Importantly, the central players of the Msps-dynein feedback loop are deeply conserved. The XMAP215 family of microtubule polymerases includes Msps in *Drosophila*, XMAP215 in *Xenopus*, ZYG-9 in *C. elegans*, Stu2/Alp14 in yeast, and ch-TOG/CKAP5 in mammals and humans (*64*). Dynein is likewise conserved as the primary minus-end motor that transports a diverse array of cargos, including mRNAs and organelles, across eukaryotes (*65–67*). Consistent with this conserved role, studies in mouse germline cysts have shown that dynein activity is required for the transfer of cytoplasmic contents, including mitochondria, Golgi, and RNAs, from sister cells into the future oocyte, a process essential for its growth and survival (*59, 61*).

Whether ch-TOG/CKAP5 contributes directly to mammalian oocyte selection remains unknown. If so, one would expect asymmetric enrichment of CKAP5 before bridge collapse, localized microtubule growth biasing plus-ends away from the future oocyte, and altered oocyte selection upon CKAP5 perturbation. Because ch-TOG is also essential for spindle assembly, future studies using temporally controlled degradation or optogenetic approaches will be required to uncouple these roles.

## Summary

In summary, our study identifies Msps/XMAP215 as a key upstream regulator of oocyte specification in *Drosophila*. We demonstrate that Msps is both necessary and sufficient for oocyte fate: loss of Msps abolishes oocyte specification, while optogenetic recruitment of Msps is sufficient to trigger it. By coupling dynein-mediated mRNA transport with microtubule polymerization, Msps establishes a positive feedback loop that ensures robust oocyte selection. These findings illustrate how germline cysts amplify a small initial asymmetry to reliably select a single oocyte from a syncytial cyst.

## Acknowledgements

We thank many colleagues who generously shared reagents: Dr Daniel St Johnston (University of Cambridge), Dr Yukiko Yamashita (Whitehead Institute, MIT), and the Bloomington *Drosophila* Stock Center (supported by NIH P40OD018537) for fly stocks; Dr Anna Akhmanova (Utrecht University) and the *Drosophila* Genomics Resource Center (DGRC, supported by NIH grant 2P40OD010949) for DNA constructs; Dr. Scott Hawley (Stowers Institute) for anti-C(3)G antibody. The anti-Orb4H8 monoclonal antibody deposited by the Paul D. Schedl group at Princeton University and the anti-Hts1B1 monoclonal antibody deposited by the Howard D. Lipshitz group at Hospital for Sick Children were obtained from the Developmental Studies Hybridoma Bank, created by the NICHD of the NIH and maintained at The University of Iowa, Department of Biology, Iowa City, IA 52242. We thank all current and past members of the Gelfand laboratory for their support, discussion, and suggestions. This study was supported by the National Institute of General Medical Sciences (2R35GM131752 to V.I.G. and R35GM145340 to G.B.G.) and the CCBx Research Program of the Center for Computational Biology of the Flatiron Institute by the Simons Foundation to V.I.G. and W.L..

## Author contributions

W.L. and V.I.G. conceived and supervised the study. W.L., G.B.G., and V.I.G. designed the experiments. W.L., M.L., H.N., and V.I.G. performed the experiments. W.L., M.L., H.N., and G.B.G. analyzed the data. W.L. and V.I.G. wrote the manuscript.

## Materials and Methods

### *Drosophila* Husbandry and Maintenance

Fly stocks and crosses were maintained on standard cornmeal food (Nutri-Fly Bloomington Formulation, Genesee, Cat. #66-121) supplemented with active dry yeast at 25 °C, except for temperature-sensitive experiments, which were conducted at the specific temperatures indicated.

The following flies were used in this study: *UAS-msps-RNAi* (HMS01906, attP40, II, Bloomington *Drosophila* Stock Center (BDSC) # 38990, targeting Msps CDS 5001-5021 nt, 5’-CTGCGCGACTATGAAGAAATA-3’) (*27*); *nos-Gal4^[VP16]^* (III)(*35, 36*); *nos-Gal4^[noVP16]^* (II); *tub-Gal80[ts]* (III) (from Dr. Yukiko Yamashita, Whitehead Institute, MIT); *osk-Gal4^[VP16]^* (III, BDSC # 44242); *Orb^[MI04761]^-GFP* (BDSC # 59817) and *MiMIC Orb^[MI04761]^* (BDSC # 37978) (*68*); *Patronin-YFP* (II, CRISPR knockin) (*69*), *Patronin-mKate* (II, CIRSPR knockin), *ubi-Patronin.RA-GFP* (X), *FRT G13 patr^[C9-C5]^* (II), and *UASp-EB1-RFP* (III, attP2) (*18*) (from Dr Daniel St Johnston, University of Cambridge); *Tub-PBac* (BDSC #8285); *UASt-NbVHH05-EGFP* (attP40, II, BDSC # 94008) (*40*); *hs-FLP^[12]^*(X, BDSC # 1929); *FRT G13* (II, BDSC #1956); *FRT G13 ubi-GFP.nls(2R1) ubi-GFP.nls(2R2)* (II, BDSC # 5826); *UASp-sfGFP.NLS* (III) (*70*); *orb^[F343]^* (BDSC # 58477)(*71, 72*).

Transgenic lines generated in this study include: *UASp-NbVHH05-3XTagRFP* (*attP14*, 36A10, PhiC31-mediated integration); *UASp-EB3N-VVDfast-mClover3-iLID* (*attP14*, 36A10, PhiC31-mediated integration); *UASp-NbVHH05-GFP-SspB* (VK22, 57F5, PhiC31-mediated integration); *Msps-EGFP-AID* (CRISPR knock-in); *Orb-TagRFP-3XHA* (MiMIC insertion), all generated by BestGene Inc.

### Plasmid Constructs

#### -pUASp-attB-NbVHH05-3XTagRFP

NbVHH05 (the alpaca nanobody against VHH05 nanotag) was PCR-amplified from pAW-NbVHH05-GFP (Addgene Plasmid #171570)(*40*) and subcloned into pUASp-attB-EMTB-3XTagRFP (*73*) via EcoRI(5’)/SpeI(3’), replacing EMTB with NbVHH05.

#### -pUASp-attB-EB3N-VVDfast-mClover3-iLID

EB3N-VVDfast-mClover3-iLID was PCR-amplified from pB80-EB3(1-189)-VVDfast-mClover3-iLID (a gift from Dr. Anna Akhmanova, Utrecht University) (*41*) and cloned into pUASp-attB via SpeI(5’)/PspXI(3’).

#### -pUASp-attB-NbVHH05-GFP-SspB

GFP-SspB was subcloned from pUASp-GFP-SspB (*73*) to replace 3XTagRFP in pUASp-attB-NbVHH05-3XTagRFP via SpeI(5’)/BamHI(3’)

#### -pUASp-Msps-3XTagRFP-3XVHH05

The 3XVHH05 tag was PCR-amplified from pScarlessHD-3XVHH05-6XMS2-DsRed (*27*) and inserted into pUASp-attB-3XTagRFP (*73*) via BamHI(5’)/XbaI(3’). An RNAi-resistant Msps (with silent mutations in the *msps-RNAi* target sequence), Msps(SM), was subcloned from pUASp-attB-Msps(SM)-YFP-BLID-K10.TLS (*27*) into pUASp-attB-3XTagRFP-3XVHH05 via NotI(5’)/SpeI(3’).

#### -pScarlessHD-5′ *msps* homology arm-C-EGFP-AID-DsRed-3′ *msps* homology arm

the construct was generated by Epoch Life Science by inserting the 5′ *msps* homology arm, EGFP, miniIAA7 (AID), DsRed, and the 3′ *msps* homology arm in the pScarlessHD-C by infusion cloning.

### CRISPR Knock-In to Create Msps-EGFP-AID

The plasmid pScarlessHD-5′ *msps* homology arm-C-EGFP-AID-DsRed-3′ *msps* homology arm was co-injected with two previously described gRNAs cloned in the pCFD5 vector (gRNA #1: GGGGTATTTCAATCAGAAGC; gRNA #2: ACGGGAAGCGCACAGTTTAT) (*27*) into *yw; nos-Cas9 attP40/CyO* flies by BestGene Inc. Progeny carrying the knock-in were identified by DsRed eye fluorescence and subsequently crossed to *Tub-PBac* flies to excise the DsRed marker via PBac transposase.

### Generation of Orb-TagRFP-3XHA Protein Trap Line via Minos Mediated Integration Cassette (MiMIC) and Recombinase Mediated Cassette Exchange (RMCE)

A protein-trap cassette intron DNA construct (DGRC, #1301, pBS-KS-attB1-2-PT-SA-SD-0-TagRFP-T-3xHA, phase 0) was injected into the MiMIC line *Orb[MI04761]* (BDSC # 37978) (*44*) in the presence of phiC31 integrase by BestGene Inc. (Supplemental Figure 4B). Transformants were identified by loss of body pigmentation (*yellow+*), and correct cassette orientation was confirmed by TagRFP expression in the ovary.

### Immunostaining in *Drosophila* Egg Chambers

Immunostaining was performed using a standard fixation and staining protocol as previously described (*70, 73, 74*). Young adult females were mated with males and fed active dry yeast for 16–18 h before dissection.

Primary antibodies used in this study: mouse monoclonal anti-Orb antibody (Orb 4H8, Developmental Studies Hybridoma Bank/DSHB, supernatant, 1:5) (*71*); mouse anti-C(3)G antibody (1A8-1G2, 1:500) (*75, 76*); mouse monoclonal anti-Hts antibody (1B1, DSHB, 1:5) (*77*). Samples were incubated with primary antibodies at 4 °C overnight. Secondary antibodies [FITC-conjugated, TRITC-conjugated, Alexa Fluor 647-conjugated anti-mouse IgG (Jackson ImmunoResearch Laboratories, Inc.; Cat# 115-095-062, Cat# 115-025-003, and Cat# 715-605-150)] were used at 10 µg/mL at room temperature (24 – 25 °C) for 4 h. In some experiments, rhodamine- or Alexa Fluor 633-conjugated phalloidin (0.2 µg/mL) and DAPI (1 µg/mL) were included. Samples were mounted in Mowiol mounting medium and dried for two days before imaging.

### Single-Molecule Inexpensive Fluorescence *in situ* Hybridization (smiFISH)

smiFISH was performed following standard protocols described previously (*27, 38*). Twenty-five 20-nucleotide-long DNA probes complementary to the 3’UTR of *msps* mRNA were designed, each carrying a 3′ FLAP-X sequence (5′-CCTCCTAAGTTTCGAGCTGGACTCAGTG-3′). Probes were mixed at equal molar ratios, and annealed with a fluorescently labeled FLAP-X probe carrying 5′ and 3′ Cy5 modifications (/5Cy5/CACTGAGTCCAGCTCGAAACTTAGGAGG-/3Cy5Sp/).

### Germline FRT Clone Induction

Virgin females of genotype *FRTG13 ubi-GFP.nls*/*CyO* or *FRTG13 ubi-GFP.nls*/*CyO; Msps-EGFP* were crossed with males carrying *hs-flp^12^*/*Y; FRTG13 patr^[C9-C5]^*/*CyO or hs-flp^12^*/*Y; FRTG13*/*CyO*. From these crosses, young pupae at 7–8 days after egg laying (AEL) were heat shocked at 37°C for 2 hours on two consecutive days. Non-CyO F1 females were collected 3-4 days after heat shock, fattened with dry active yeast overnight, and dissected for Orb staining.

### Live Imaging of *Drosophila* Egg Chambers

Young female adults were mated with males and fed active dry yeast for 16 to 18 h before dissection. Ovaries were dissected in Halocarbon oil 700 (Sigma-Aldrich, Cat# H8898) as described previously (*12, 13, 27, 70, 73, 74*). Freshly dissected samples were imaged on a Nikon W1 spinning disk confocal microscope (Yokogawa CSU, pinhole size 50 µm) equipped with a Hamamatsu ORCA-Fusion digital CMOS camera and a 40×/1.25 N.A. silicone oil lens, controlled by Nikon Elements software.

### OptoMsps Recruitment in *Drosophila* S2R+ Cells

*Drosophila* S2R+ cells (DGRC Stock #150; RRID: CVCL_Z831) were maintained in Insect-Xpress medium (Lonza, Catalog # 12-730Q) in a 25°C incubator and were transiently transfected with pAC-Gal4, pUASp-EB3N-VVDfast-mClover3-iLID, pUASp-NbVHH05-GFP-SspB, and pUASp-Msps-3XTagRFP-3XVHH05 at a 4:3:3:1 ratio using Effectene (Qiagen, Cat# 301425) following the manufacturer’s instructions. After 2-3 days, cells were plated on Concanavalin A (ConA)-coated coverslips and imaged on a Nikon W1 spinning disk confocal microscope (Yokogawa CSU, pinhole size 50 µm) equipped with a Hamamatsu ORCA-Fusion digital CMOS camera and a 100×/1.35 N.A. silicone oil lens, controlled by Nikon Elements software. Optogenetic recruitment was induced by a 470 nm Lumencor LED light controlled by a Mightex Polygon DMD Illuminator at 0.8% power, with 30 s of continuous illumination per cycle, followed by acquisition using a 561 nm laser at a single focal plane and an end-point final acquisition using a 488 nm laser. All the procedures were performed under red-light-only conditions to minimize unintended activation from ambient light.

### OptoMsps Recruitment in *Drosophila* Ovaries with EB1-RFP

Recombination between (1) *UASp-EB3N-VVDfast-mClover3-iLID* (attP14) and *UASp-NbVHH05-GFP-SspB (VK22)*; (2) *osk-Gal4[VP16]* (III) and *Msps-3XVHH05-6XMS2* (89B1-89B2); (3) *UASp-EB1-RFP* (attP2) and *Msps-3XVHH05-6XMS2* (89B1-89B2) were performed following standard protocols. F1 females (*yw*; *UASp-EB3N-VVDfast-mClover3-iLID, UASp-NbVHH05-GFP-SspB*/*CyO; osk-Gal4[VP16], Msps-3XVHH05-6XMS2/UASp-EB1-RFP, Msps-3XVHH05-6XMS2*) were dissected in Halocarbon oil 700 and imaged on a Nikon W1 spinning disk confocal microscope (Yokogawa CSU, pinhole size 50 µm) equipped with a Hamamatsu ORCA-Fusion digital CMOS camera and a 40×/1.25 N.A. silicone oil lens, controlled by Nikon Elements software. Optogenetic recruitment was induced by a 470 nm Lumencor LED light controlled by a Mightex Polygon DMD Illuminator at 0.8% power, with 30 s continuous illumination per cycle. Imaging was performed using a 561 nm laser at a single focal plane 2 s intervals for a total of 10 s. For the results (Figure 4), only the first frame of each 10-s time lapse per illumination cycle was shown and analyzed; sum projections of the full 10-s time lapses are presented in Video 7. All procedures were performed under red-light-only conditions to prevent unintended activation by ambient light.

### OptoMsps Recruitment in *Drosophila* Ovaries with Orb-TagRFP

Recombination between *Msps-3XVHH05-6XMS2* (89B1-89B2) and *Orb-TagRFP* (94E9) was performed following standard protocols. F1 females of *yw*; *UASp-EB3N-VVDfast-mClover3-iLID, UASp-NbVHH05-GFP-SspB*/*CyO; osk-Gal4[VP16], Msps-3XVHH05-6XMS2/UASp-EB1-RFP, Msps-3XVHH05-6XMS2* were dissected in supplemented Schneider’s medium (pH 6.9; 20 mM HEPES, 0.5 mg/mL Bovine Serum Albumin (BSA), 5mM Glucose, 20% fetal bovine serum (FBS), 5 μg/ml insulin, 100 μg/ml penicillin-streptomycin, 10 μg/ml tetracycline, and 5 μM 20-Hydroxyecdysone) containing 25 μM biliverdin dihydrochloride (MP Biomedicals Cat #194886) for far-red membrane labeling. Ovarioles in supplemented Schneider’s medium were mixed with BME hydrogel (R&D Systems, BME001-01) at a volume ratio of 2:1 and mounted in a “sandwich” configuration between a lummox membrane (Sarstedt, # 94.6150.101) and a glass coverslip, using acid-washed ≤106 μm (−140 U.S. sieve) glass beads (Sigma Aldrich, G4649-10G) as the spacer and silicone vacuum grease (Beckman, 335148) as the glue. Samples were allowed to briefly solidify at 30 °C for 15–20min before imaging. Imaging was performed on a Nikon W1 spinning disk confocal microscope (Yokogawa CSU, pinhole size 50 µm) equipped with a Hamamatsu ORCA-Fusion digital CMOS camera and a 40×/1.25 N.A. silicone oil lens, controlled by Nikon Elements software.

Local optogenetic recruitment was induced by a 470 nm Lumencor LED light controlled by a Mightex Polygon DMD Illuminator at 0.8% power and continuous illumination for 2 min per cycle, followed by acquisition with 561nm and 640nm lasers over an 18-µm z-stack (3 µm each step, a total of 7 steps). Sum slices of the z-stacks were used for analysis and presentation (Figure 4; Video 8). Global optogenetic recruitment was induced by acquisition using a 488 nm laser at 2s intervals for 2 min at a single focal plane (Video 6). All procedures were performed under red-light-only conditions to avoid unintended activation by ambient light.

### Egl-mRNA binding assay

The *in vivo* RNA binding experiment was performed as previously described (*47, 78*). In brief, ovarian lysates were prepared from wild-type flies. 600 µg of total lysate was used in each immunoprecipitation, using either an anti-HA antibody (control) or an antibody against Egl. The co-precipitating RNAs were eluted, extracted, and reverse transcribed using Superscript III (Life Technologies). SsoAdvanced Universal SYBR green supermix (BioRad) was used to perform the quantitative PCR reaction using a Bio-Rad CFX96 Real-time PCR machine. Fold enrichment was calculated by normalizing the amount of *msps*, *nos*, and *grk* mRNAs that precipitated with the control anti-HA antibody versus the anti-Egl antibody.

### Statistical Analysis

Figures display either percentages of phenotypes or mean values, as indicated in the corresponding legends. Error bars represent SEM, and N represents the number of samples analyzed per assay. Statistical significance was assigned as follows: P ≥ 0.05, not significant (n.s.); 0.01 ≤ P < 0.05, significant (*); 0.001≤ P < 0.01, very significant (**); P 0.0001 ≤ P < 0.001, extremely significant (***); P < 0.0001, extremely significant (****).

## Supplementary Figures and Supplementary Figure Legends

**Supplementary Figure 1.**
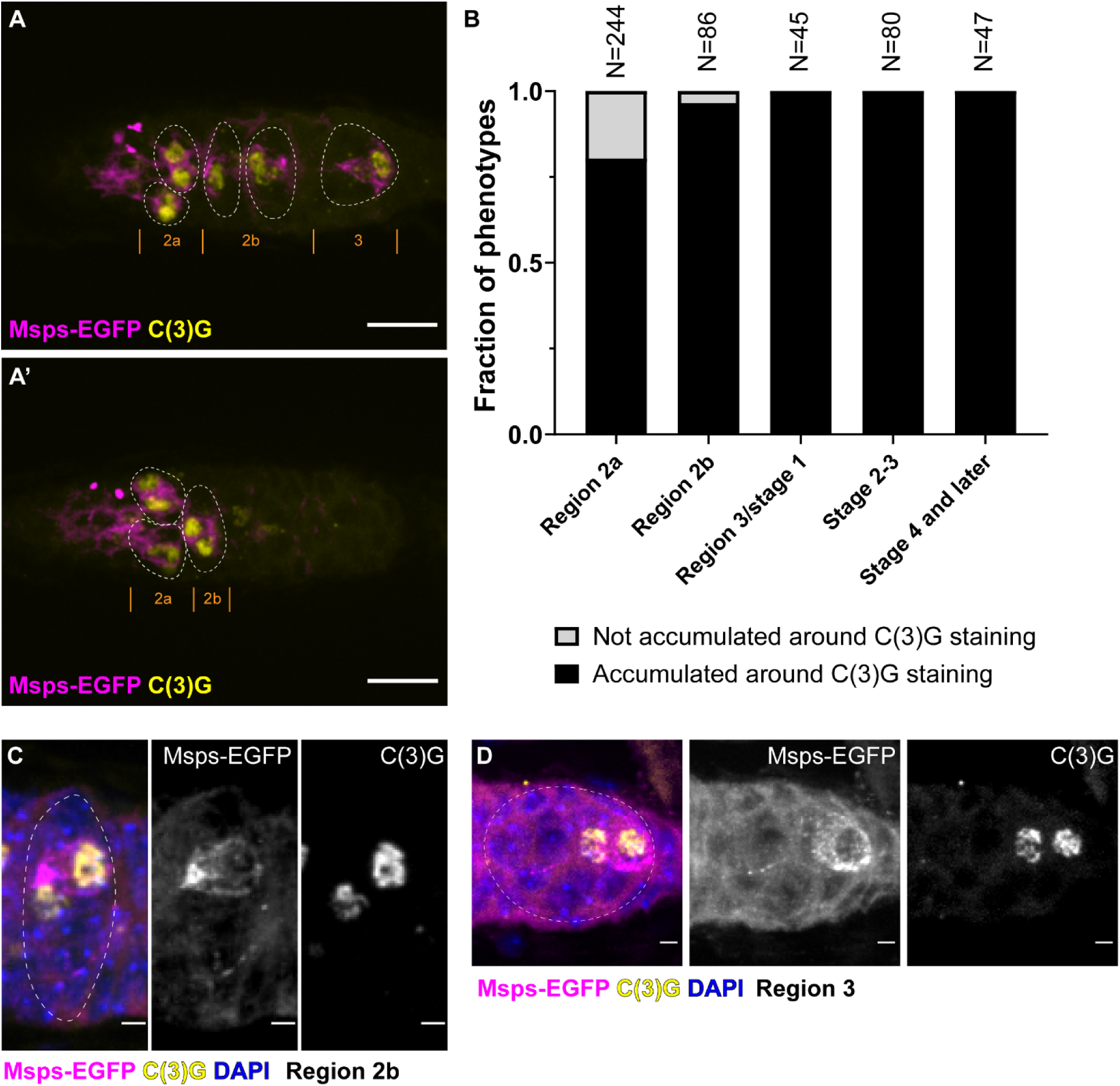
Msps protein expression in pro-oocytes and early oocytes. Related to Figure 1. (A–A’) Representative germarium showing Msps-EGFP localization and staining of the meiotic marker C(3)G from region 2a to region 3. Two focal planes from the same sample are shown. Scale bars, 10 µm. (B) Quantification of the percentage of oocytes with Msps-EGFP enrichment around C(3)G staining at different stages. (C–D) Examples of two pro-oocytes both in meiosis, but only one showing strong Msps enrichment, in region 2b (C) and region 3 (D). Scale bars, 2 µm.

**Supplementary Figure 2.**
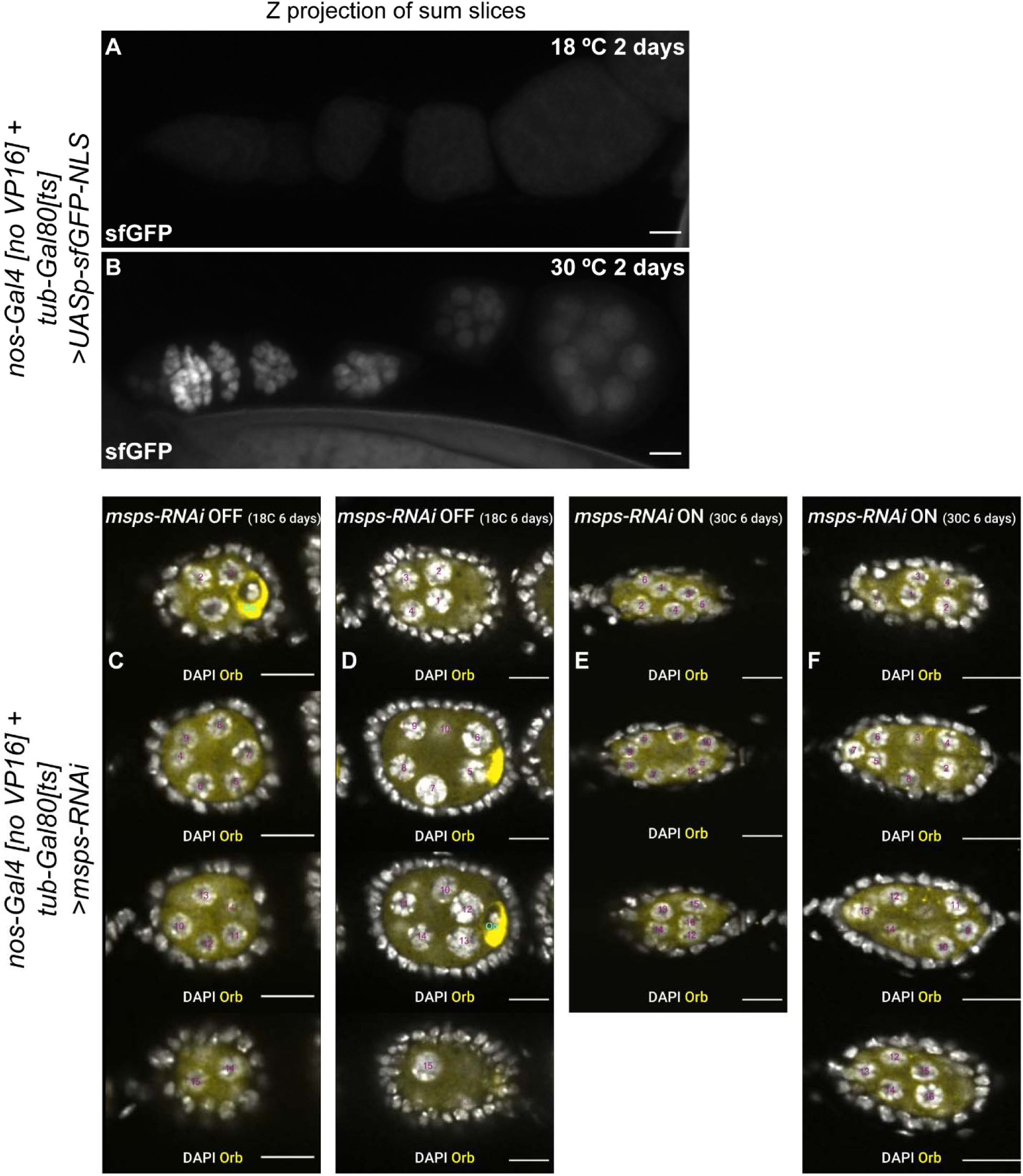
Temperature shift controls germline expression, and loss of Msps abolishes oocyte specification. Related to Figure 2. (A–B) Germline expression of nuclear-localized sfGFP under the temperature-sensitive Gal80 system. Z-projected sum slices are shown for samples at 18 °C and 30 °C. (C–F) Germline cell number counts in *msps-RNAi* OFF (C–D) and ON (E–F) samples. The 15 polyploid nurse cells are numbered in purple, and the diploid oocyte with Orb accumulation is labeled “Oo” in cyan. See also Videos 2–3. (A–F) Scale bars, 10 µm.

**Supplementary Figure 3.**
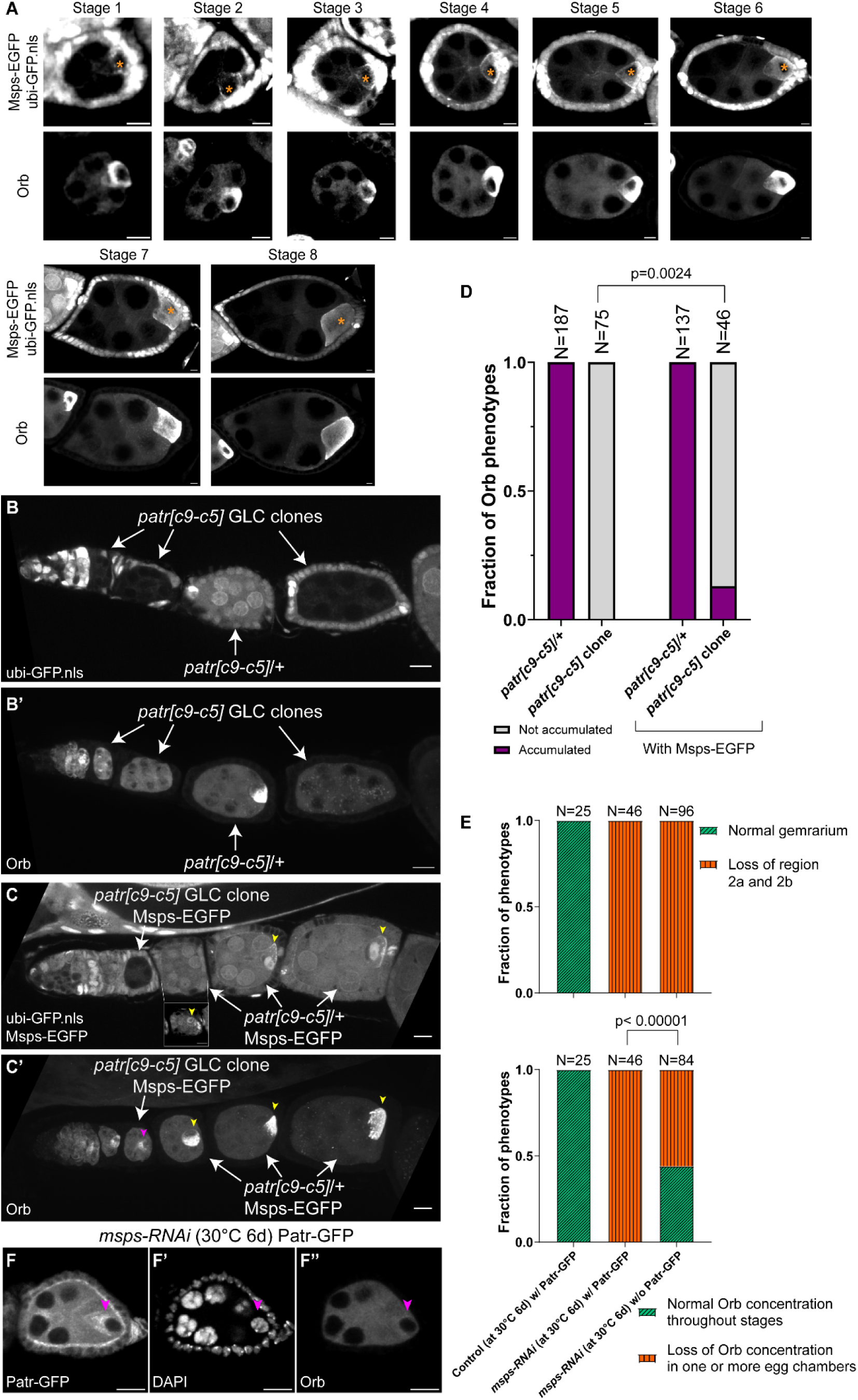
Genetic interaction between Msps and Patronin. Related to Figure 3. (A) Msps-EGFP localization (top) and Orb staining (bottom) in FRT G13 control clones from stage 1 to stage 8. The absence of the clonal marker, nuclear GFP (GFP.nls), indicates the presence of germline clones. Orange asterisks mark the oocytes. Scale bars, 5 µm. (B-B’) Orb staining in *patr[c9-c5]* heterozygous and homozygous egg chambers. *patr[c9-c5]* germline clones (GLC) are identified by the absence of nuclear GFP (GFP.nls). Scale bars, 10 µm. (C-C’) Orb staining in *patr[c9-c5]* heterozygous and homozygous egg chambers with one endogenous copy of Msps tagged with EGFP (Msps-EGFP). (C) Yellow arrowheads, Msps-EGFP accumulation in the oocyte alongside the clonal marker (nuclear GFP); inset, Msps-EGFP accumulation in the oocyte at a different focal plane. (C’) Magenta arrowhead, Orb enrichment in the *patr[c9-c5]* homozygous clone. (D) Quantification of Orb phenotypes in *patr[c9-c5]* heterozygous and homozygous egg chambers, with or without Msps-EGFP. (E) Quantification of germaria lacking region 2a/2b and of oocyte specification defects in controls and *msps-RNAi* with or without ubi-Patronin-GFP. Statistical comparisons between the two groups were performed using Fisher’s exact test. Note: the *msps-RNAi* (30 °C, 6 days) dataset without Patr-GFP is the same as in Figure 2F.

**Supplementary Figure 4.**
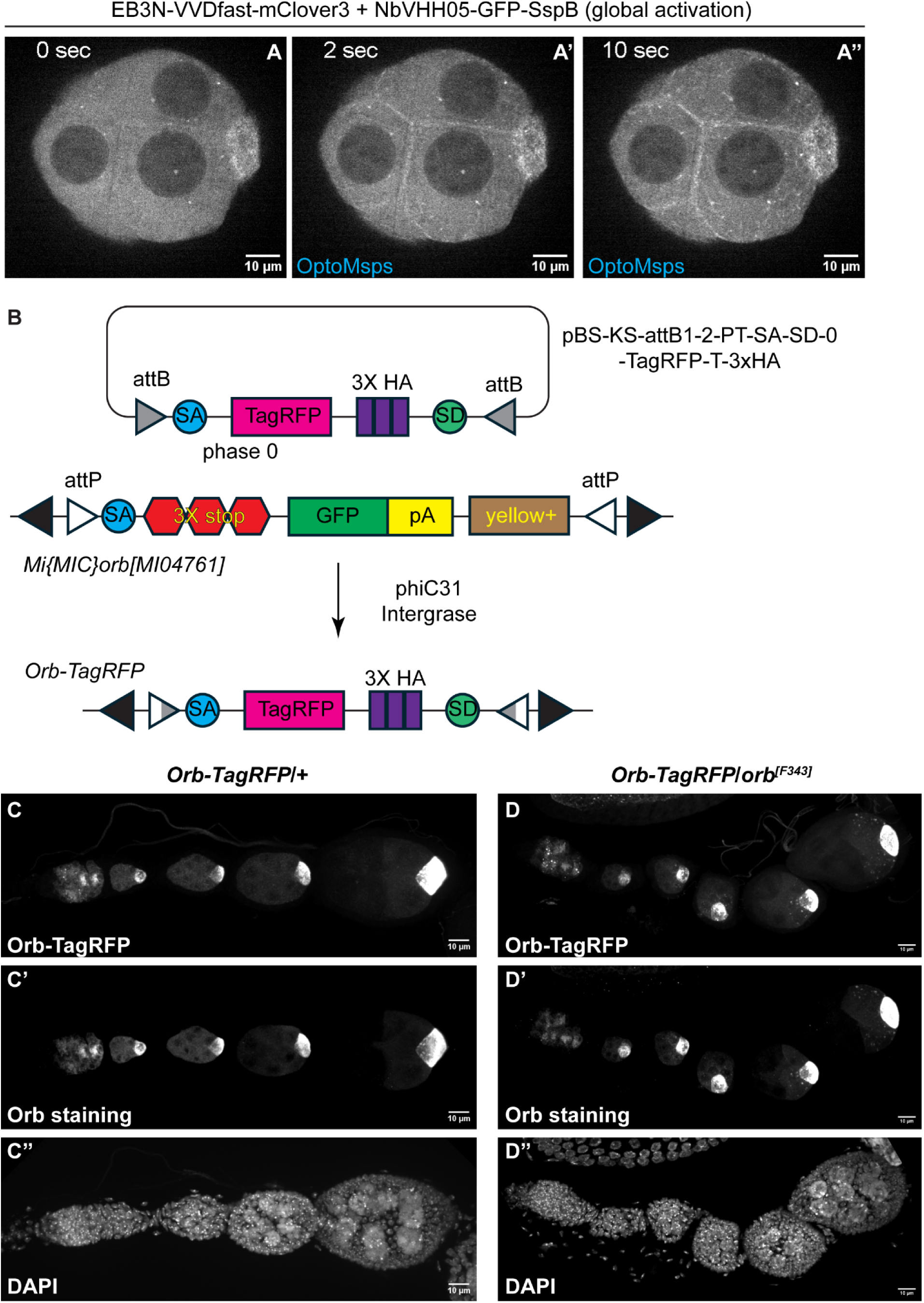
Global activation of OptoMsps and generation of the Orb-TagRFP line. Related to Figure 4. (A–A’’) Global activation of OptoMsps in a *Drosophila* egg chamber. Global activation was achieved via image acquisition using a 488 nm laser. See also Video 6. (B) Schematic of the Orb-TagRFP line generated with the MiMiC system. A DNA cassette containing TagRFP-3×HA (phase 0) was injected into the *Orb[MI04761]* line in the presence of PhiC31 integrase. Recombination between attP and attB sites generates an endogenously tagged Orb-TagRFP line (see Materials and Methods for more details). (C–C’’) Representative images of Orb staining in an Orb-TagRFP/+ ovariole. Orb antibody staining and Orb-TagRFP fluorescence colocalize throughout oogenesis. (D-D’’) Representative images of Orb staining in an Orb-TagRFP/*orb[F343]* ovariole. *orb[F343]* is an amorphic allele of *orb*, with premature termination at Q240t. Orb antibody staining and Orb-TagRFP fluorescence colocalize throughout oogenesis. *Orb-TagRFP/orb[F343]* females undergo normal oogenesis, produce eggs of normal size, and are fully fertile.

**Supplementary Figure 5.**
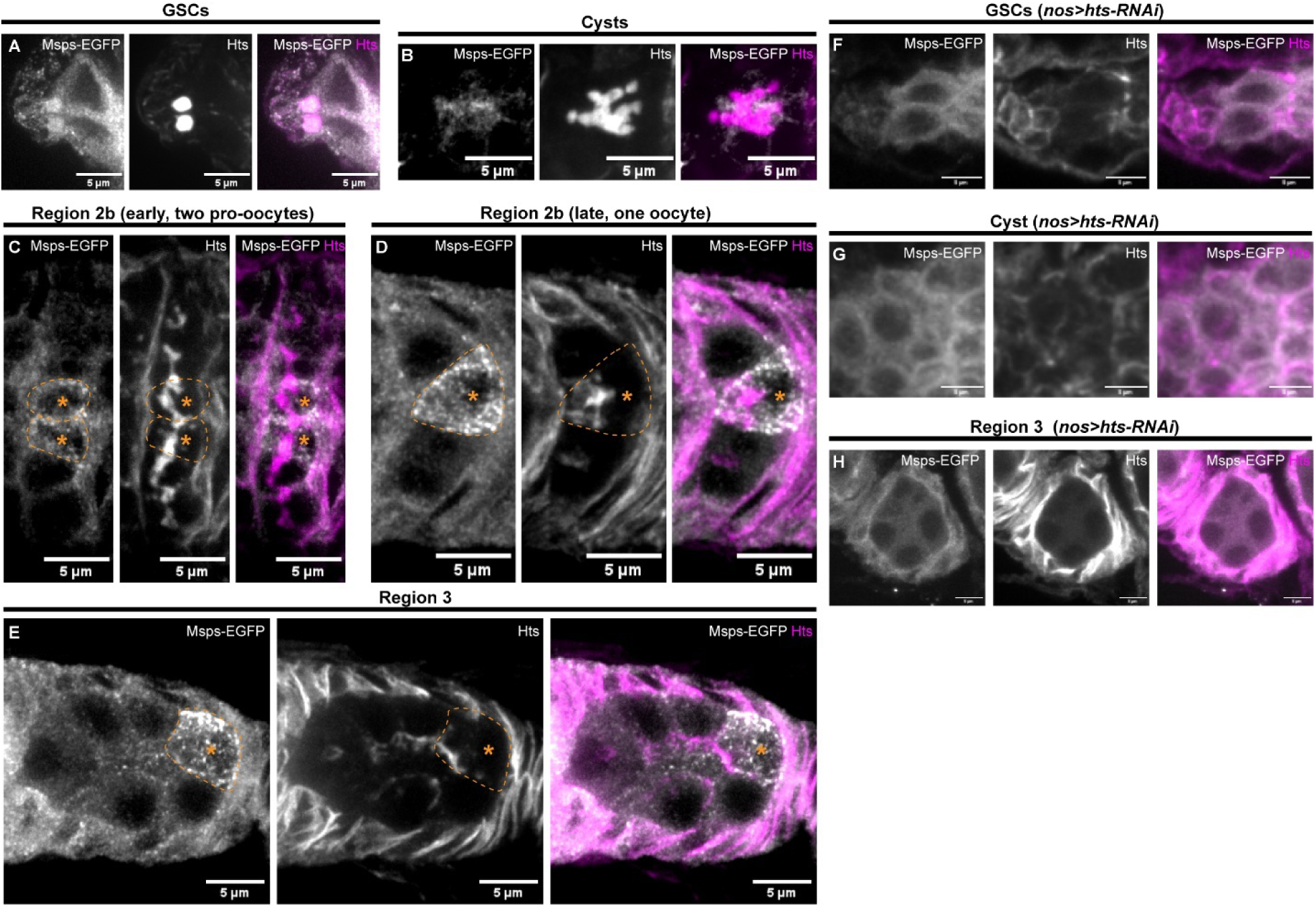
Msps localization to spectrosome/fusome structures and asymmetric segregation into pro-oocytes and the oocyte. Related to Figure 5. (A–E) Msps-EGFP localization with Hts staining at different stages of the germarium. (A) Msps is enriched at the spectrosomes in GSCs. (B) Msps shows a looser association with the fusome in cysts. (C–E) Msps-EGFP becomes concentrated in pro-oocytes (C) and the presumptive oocyte (D–E), losing its clear association with fusome structures. (F–H) Msps-EGFP localization in *hts-RNAi* knockdown samples. No spectrosome or fusome association of Msps-EGFP is detected in GSCs (F) or cysts (G), and Msps-EGFP fails to accumulate in region 3 (H). Note that *hts-RNAi* disrupts germline cell divisions, resulting in fewer than 16 germline cells per cyst.

## Supplementary Video Legends

**Video 1. Msps-EGFP dynamics in pro-oocytes and oocytes of the *Drosophila* germarium.** Two-minute time-lapse imaging of the CRISPR knock-in line Msps-EGFP shows Msps-EGFP–positive comets in regions 2a, 2b, and 3. Related to Figure 1.

**Video 2. Normal oocyte specification in *msps-RNAi* OFF samples.** Two representative egg chambers from flies carrying germline *msps-RNAi* under the temperature-sensitive Gal80 system, raised and maintained at 18 °C for 6 days (RNAi OFF). Both chambers contain 15 polyploid nurse cells (numbered in purple) and one diploid oocyte with proper Orb accumulation (labeled as “Oo” in cyan). Scale bar, 10 µm. Related to Figure 2.

**Video 3. No oocyte specification in *msps-RNAi* ON samples.** Two representative egg chambers from flies carrying germline *msps-RNAi* under the temperature-sensitive Gal80 system, raised at 18 °C and shifted to 30 °C for 6 days (RNAi ON). Both chambers contain 16 polyploid nurse cells (numbered in purple) with no oocyte or Orb enrichment. Scale bar, 10 µm. Related to Figure 2.

**Video 4. Dual labeling of Msps and Patronin in the germarium shows their simultaneous accumulation in the early oocyte.** EGFP-tagged Msps (left), mKate-tagged Patronin (middle), and merged channel (right). Both examples are from the same germarium at different focal planes. Related to Figure 3.

**Video 5. Local recruitment of Msps using OptoMsps in *Drosophila* S2R+ cells.** Blue-light illumination (blue box) induced recruitment of Msps-3XTagRFP-3XVHH05 to microtubules. Related to Figure 4. See also “OptoMsps recruitment in *Drosophila* S2R+ cells” in Materials and Methods for more details.

**Video 6. Global recruitment of OptoMsps in a representative *Drosophila* egg chamber.** Global activation was triggered by green-channel acquisition of EB3N-VVDfast-mClover3 and NbVHH05-GFP-SspB, with clear recruitment onto microtubules visible within 2 seconds of illumination. Related to Figure 4.

**Video 7. OptoMsps locally increases EB1-RFP signal in *Drosophila* nurse cells.** Blue-light illumination (blue box) induced an elevated EB1-RFP signal, which subsequently spread throughout the entire nurse cell over time. Related to Figure 4. See also “OptoMsps recruitment in *Drosophila* ovaries with EB1-RFP” in Materials and Methods.

**Video 8. OptoMsps recruitment elevates Orb-TagRFP signal in *Drosophila* nurse cells.** Blue-light–induced recruitment of Msps (blue box) led to an increase in Orb-TagRFP within nurse cells. Related to Figure 4. See also “OptoMsps recruitment in Drosophila ovaries with Orb-TagRFP” in Materials and Methods.

